# Genome-wide profiling of histone H3K4me3 and H3K27me3 modifications in individual blastocysts by CUT&Tag without a solid support (NON-TiE-UP CUT&Tag)

**DOI:** 10.1101/2022.02.07.479478

**Authors:** Kazuki Susami, Shuntaro Ikeda, Yoichiro Hoshino, Shinnosuke Honda, Naojiro Minami

**Affiliations:** Laboratory of Reproductive Biology, Graduate School of Agriculture, Kyoto University, Kyoto 606-8502, Japan

## Abstract

Individual analysis of the epigenome of preimplantation embryos is useful for characterizing each embryo and for investigating the effects of environmental factors on their epigenome. However, it is difficult to analyze genome-wide epigenetic modifications, especially histone modifications, in a large number of single embryos due to the small number of cells and the complexity of the analysis methods. To solve this problem, we further modified the CUT&Tag method, which can analyze histone modifications in a small number of cells, such that the embryo is handled as a cell mass in the reaction solutions in the absence of the solid-phase magnetic beads that are used for antibody and enzyme reactions in the original method (NON-TiE-UP CUT&Tag; NTU-CAT). By using bovine blastocysts as a model, we showed that genome-wide profiles of representative histone modifications, H3K4me3 and H3K27me3, could be obtained by NTU-CAT that are in overall agreement with the conventional chromatin immunoprecipitation-sequencing (ChIP-seq) method, even from single embryos. However, this new approach has limitations that require attention, including false positive peaks and lower resolution for broad modifications. Despite these limitations, we conclude that NTU-CAT is a promising replacement for ChIP-seq with the great advantage of being able to analyze individual embryos.

## Introduction

A close investigation of the epigenome of early preimplantation embryos is important for validating the temporal and spatial regulation of gene expression that is crucial for their development and examining the impact of environmental factors on their epigenome. Post-translational histone modifications are one of the representative epigenetic modifications, and genome-wide investigations of these modifications in early embryos have been performed using mainly the chromatin immunoprecipitation-sequencing (ChIP-seq) method ^1-3^. However, because early embryos are composed of a very small number of cells (∼100 cells), it is difficult to analyze a single embryo. However, several methods have been proposed recently to overcome the disadvantages of ChIP-seq, which requires a large number of cells for analysis, and these methods have been applied to the analysis of single or small numbers of preimplantation embryos ^4,5^. First, Henikoff and colleagues, based on Laemmli’s Chromatin ImmunoCleavage (ChIC) strategy ^6^, proposed the Cleavage Under Targets and Release Using Nuclease (CUT&RUN) ^7,8^. Although CUT&RUN can produce high quality data from as few as 100-1000 cells, it has disadvantages in terms of the time, cost, and effort required for purification, end polishing, and adapter ligation of micrococcal nuclease (MNase)-cleaved DNA fragments that are released into the supernatant ^7,8^. However, they overcame these disadvantages and developed a new method called Cleavage Under Targets and Tagmentation (CUT&Tag), which uses a fusion protein of Tn5 transposase and protein A (pA-Tn5), instead of pA-MNase, loaded with sequencing adapters to ligate the adapters directly to the cleaved fragments ^9,10^. This strategy results in the formation of PCR-amplifiable adapter-ligated fragments for sequencing *in situ* (not released into the supernatant) unlike in the case of CUT&RUN. Therefore, we expected that CUT&Tag could be performed without binding cells or isolated cell nuclei to the solid phase, as used in original CUT&RUN and CUT&Tag. Here, we report profiling of representative histone modifications, H3K4me3 and H3K27me3, in single bovine blastocysts as a model using CUT&Tag without a solid phase (NON-TiE-UP CUT&Tag; NTU-CAT).

## Results

### Schema for NTU-CAT

The schema for NTU-CAT is shown in Fig. 1. After removal of the zona pellucida, the embryos were immersed in a primary antibody solution containing a detergent (digitonin) and incubated without binding of the cells or isolated nuclei to a solid phase (concanavalin A-coated magnetic beads) as in the conventional method. Subsequent secondary antibody reactions, tethering of pA-Tn5 fusion protein with sequencing adapters, and tagmentation were also performed without a solid support, by transferring the embryos to the respective reaction solutions. The tagmented DNA was extracted and PCR amplified using index primers, and the library after purification was used for sequencing. The numbers of sequencing reads generated and processed are summarized in Table S1.

**Figure 1.**
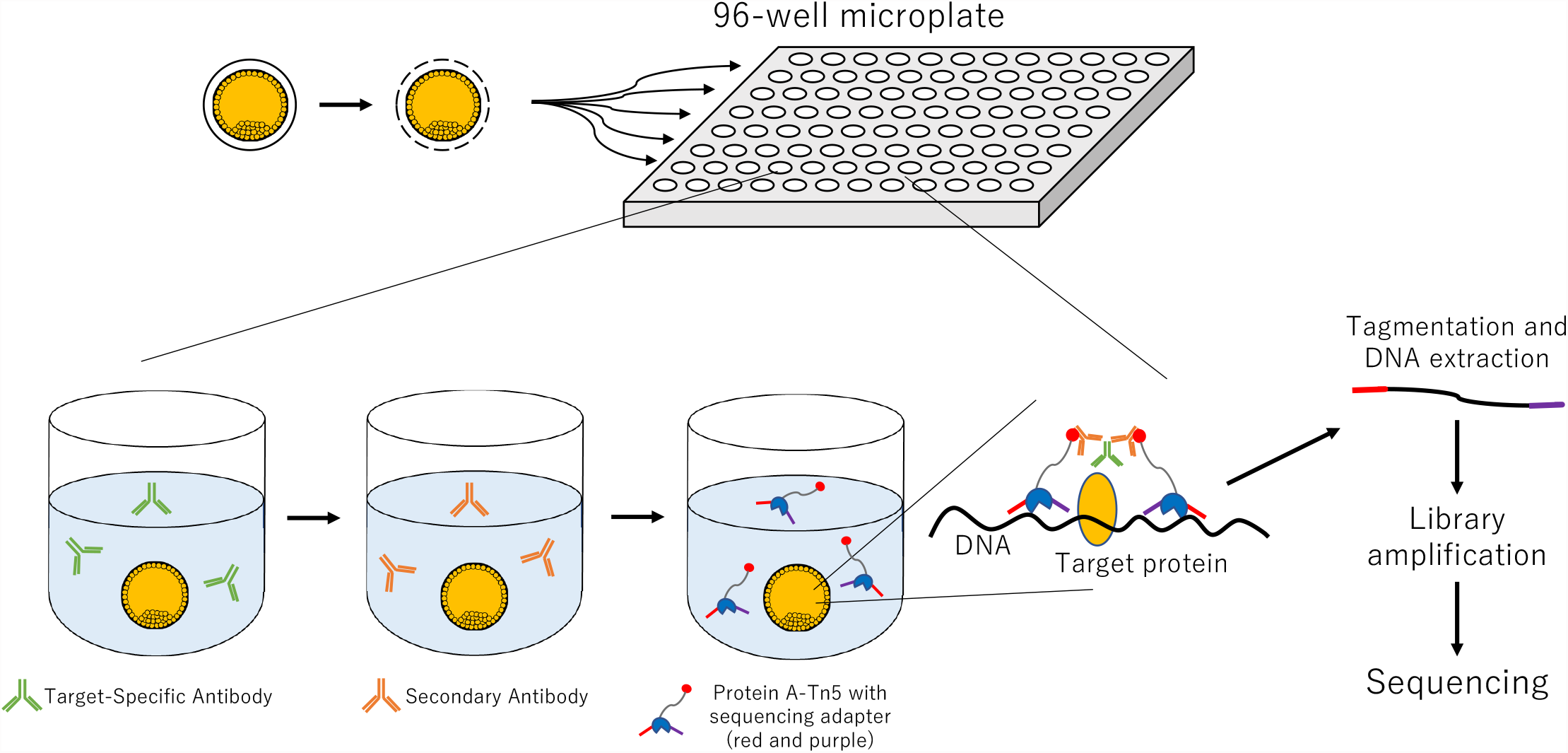
Schematic diagram of NON-TiE-UP CUT&Tag (NTU-CAT). See the Methods section for details.

### H3K4me3 profile of blastocysts assessed by NTU-CAT

Among the various types of histone methylation, H3K4me3 is a typical transcription-promoting marker that is relatively easy to analyze because it generally shows sharp and distinct modifications ^1-3^. Therefore, we initially targeted this modification for NTU-CAT application. Because NTU-CAT allows for rapid analysis of histone modifications from individual embryos, we could readily obtain data from many embryos. A snapshot of the NTU-CAT peaks from nine replicates (i.e., nine single blastocysts) is shown in Fig. 2a, alongside the peaks from our previous ChIP-seq analysis of a cohort of blastocysts for H3K4me3 modification ^11^. The overall landscape depicted by the location and shape of the peaks was very similar between both methods, except for subtle differences in the shape of the peaks (Fig. 2a). Fig. 2b shows the average profile plots of the H3K4me3 signal around the transcript start site (TSS) regions in these experiments. A striking difference between the NTU-CAT and ChIP-seq profiles was that the valley-like shapes near TSSs detected in ChIP-seq were not detected in NTU-CAT. However, the peak shape of NTU-CAT could be approximated to that of ChIP-seq. Supportively, pairwise comparisons of H3K4me3 signals showed a high correlation between the methods and replicates (Fig. 3).

**Figure 2.**
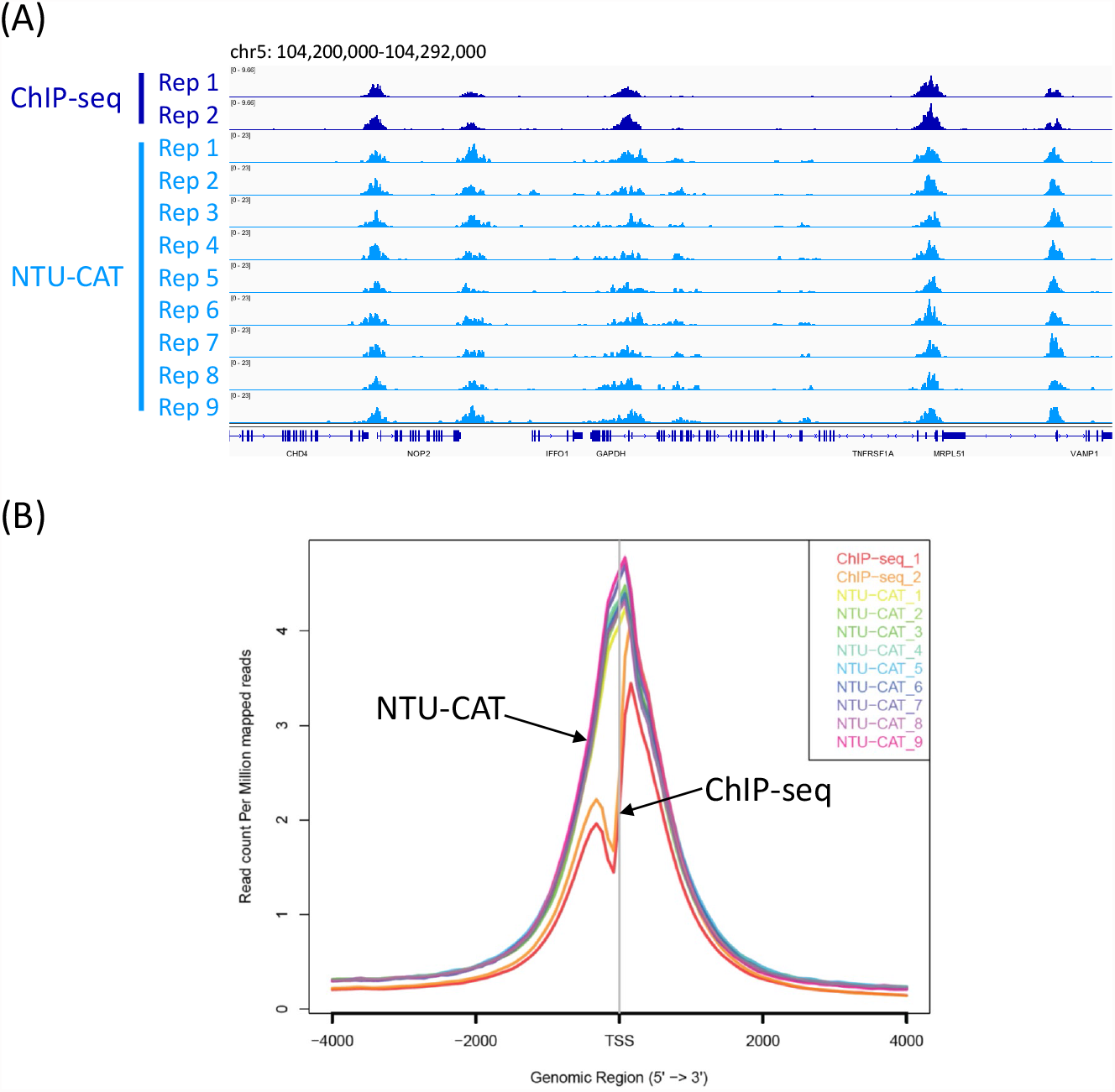
Comparison of NTU-CAT and ChIP-seq results for H3K4me3 in bovine blastocysts. (a) Snapshot of the H3K4me3 landscape in a 92-kb region (chromosome 5) that includes the *GAPDH* TSS. The NTU-CAT-derived peaks in nine replicates were visualized using the Integrative Genomics Viewer ^19^ alongside the peaks from our previous ChIP-seq analysis of a cohort of blastocysts ^11^. (b) Average profile plot of the H3K4me3 signal around genome-wide TSSs. The signals from the nine replicates of NTU-CAT and duplicates of ChIP-seq ^11^ are shown. The average profile plots were generated by ngs.plot ^18^.

**Figure 3.**
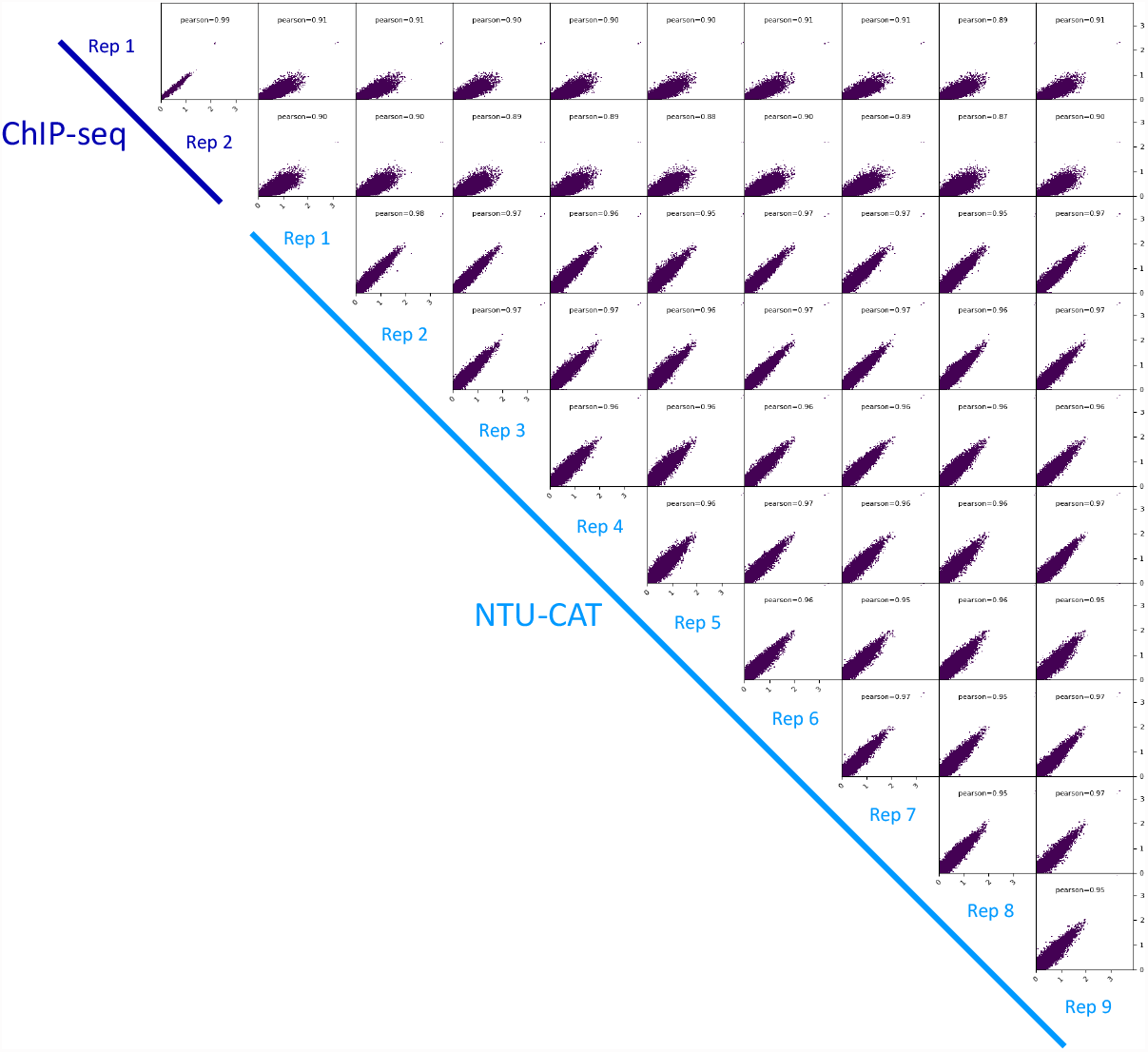
Correlation analysis of duplicates of ChIP-seq and nine replicates of NTU-CAT for H3K4me3 in bovine blastocysts. Scatterplots of these pairwise comparisons are shown with Pearson correlation coefficients. Bin sizes of 10 kb and natural log transformation after adding 1 were used for drawing with deepTools (https://deeptools.readthedocs.io/en/develop/).

We detected approximately 20,000 significant peaks throughout the genome (Figs. 4a and 5a) and 20% of the peaks were located on gene promotor regions (Fig. 4b). In addition, average profile plotting around gene bodies categorized by gene groups with different expression levels revealed that highly expressed gene groups had more extensive H3K4me3 modifications (Fig. 4c). Although these results were obtained from single embryos with a smaller number of mapped reads, they are in complete agreement with our previous results using conventional ChIP-seq with a cohort of embryos ^11^.

**Figure 4.**
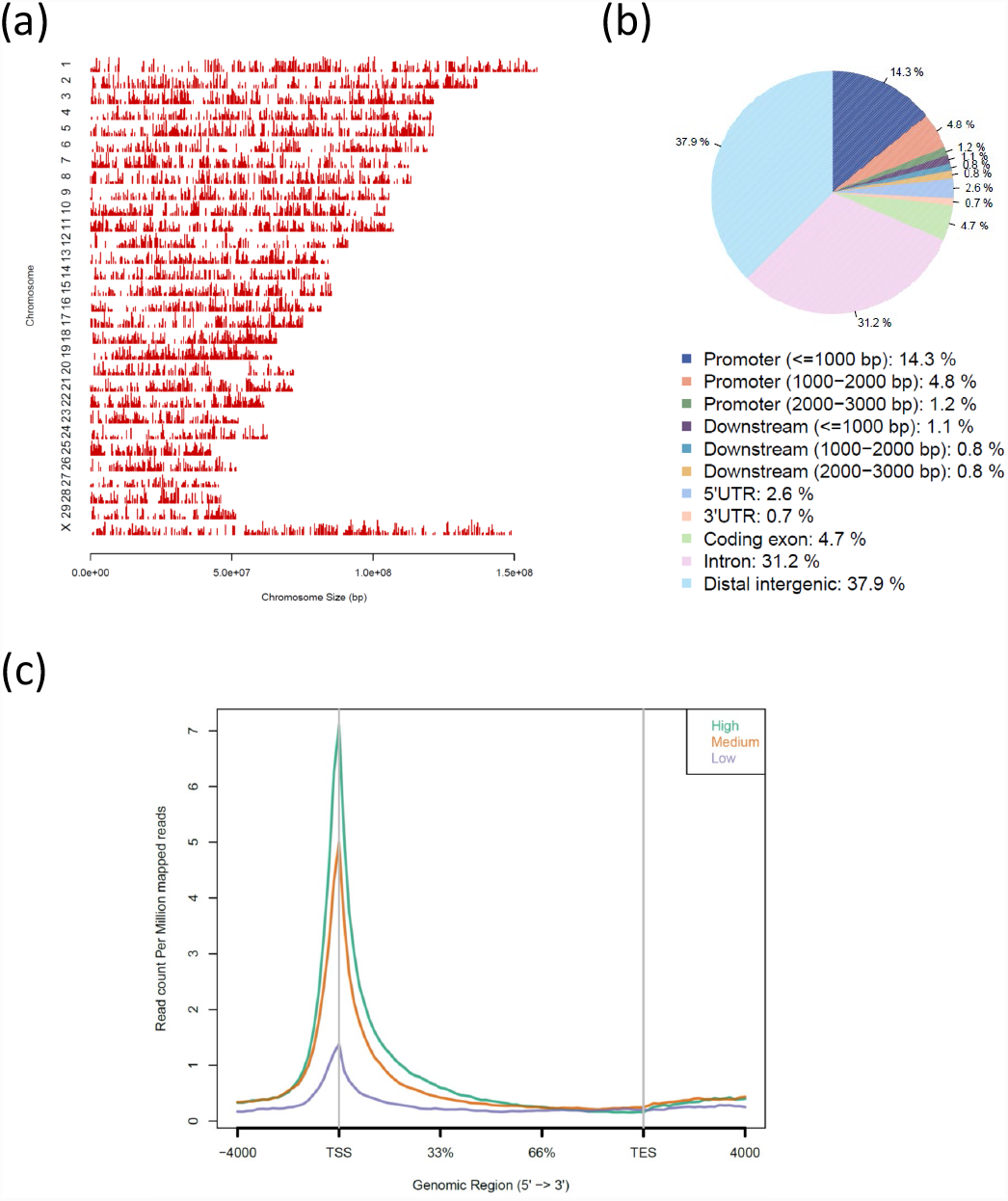
Overview of the NTU-CAT results for H3K4me3 in bovine blastocysts. (a and b) Distribution of H3K4me3 peaks in each chromosome (a) and in corresponding genic and intergenic regions (b). These figures were generated by the CEAS ^17^ tool. (c) Average profile plot around gene bodies categorized by gene groups with different expression levels based on GSE52415 ^20^. The figure was generated by ngs.plot ^18^. All of the figures were made using the Rep1 sample in Figs. 1 and 2.

CUT&Tag is a Tn5 transposase-based method ^9,10^ and there are concerns of possible bias in epigenomic profiling caused by the preference of Tn5 transposase to cut open chromatin regions ^12^. We validated this point in the following analyses. In order to identify the open chromatin regions, we used publicly available ATAC-seq data of bovine inner cell mass (GSM4516360) deposited by Halstead et al., ^13^ and mapping and peak calling generated 137,149 ATAC-seq peaks. We then calculated the false positive rate (FPR) possibly caused by the bias of Tn5 transposase toward open chromatin, where the FPR was the number of peaks that did not overlap with ChIP-seq, but did overlap with the ATAC-seq peaks, divided by the total number of peaks, as proposed by Wang et al. ^12^. As a result, the FPR rate was 10%-15% (Fig. 5a). Furthermore, the areas of the false positive peaks were relatively smaller than those of all peaks (Fig. 5b).

**Figure 5.**
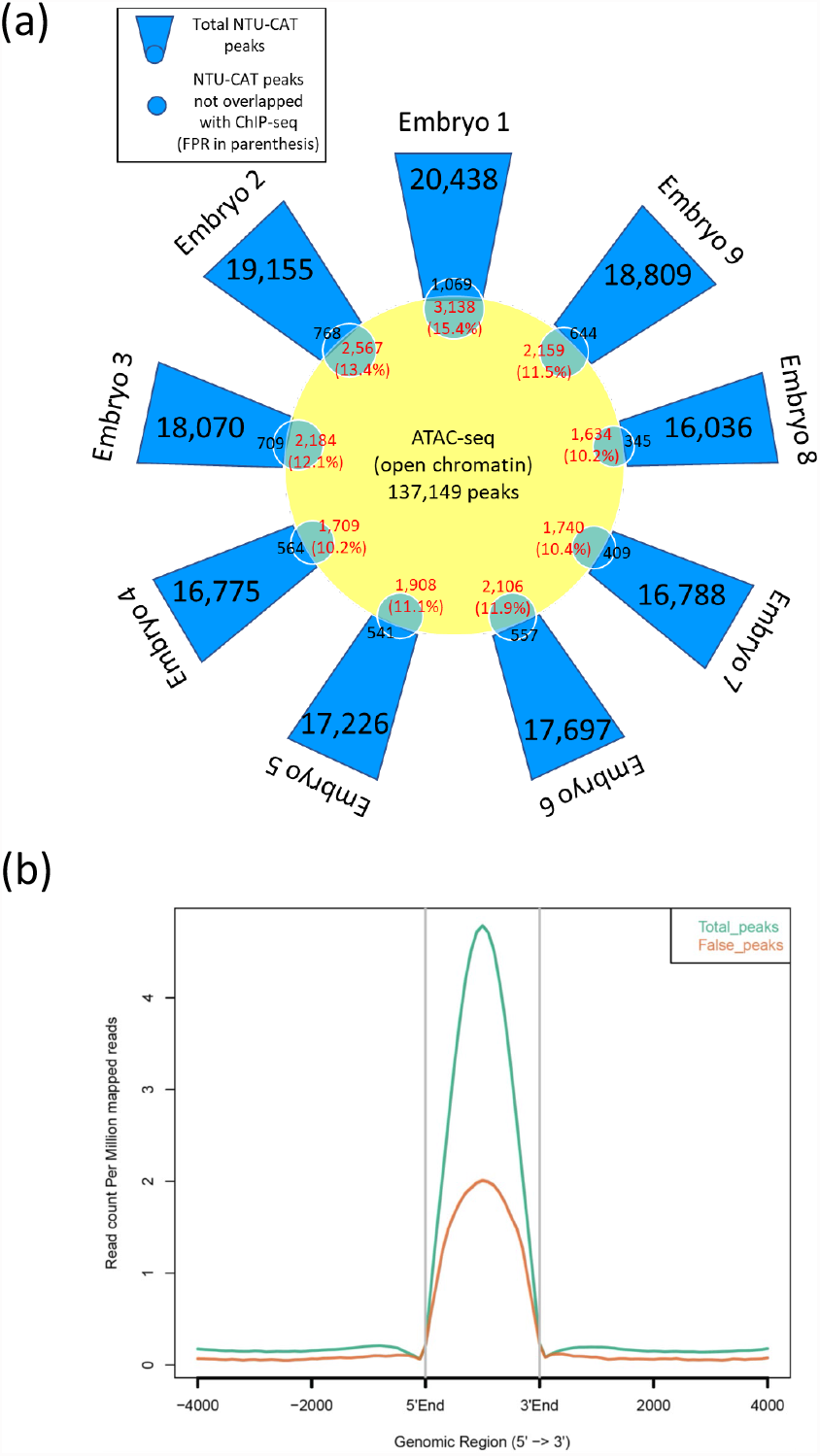
The validation of false positive peaks in NTU-CAT possibly due to the open chromatin bias of Tn5 transposase. (a) A girasol plot showing the number of peaks and false positive rate (FPR). The FPR was calculated by the number of NTU-CAT peaks that did not overlap with ChIP-seq (Blastocyst 1 of our previous report ^11^) but did overlap with the ATAC-seq peaks ^13^, divided by the total number of peaks ^12^. (b) Average profile plot of the H3K4me3 signals in total NTU-CAT peaks and false positive peaks in Rep1 (Embryo 1 in Fig. 1a). The figure was generated by ngs.plot ^18^

### H3K27me3 profile in blastocysts assessed by NTU-CAT

Compared with H3K4me3 modification, H3K27me3 is a typical transcription-repressing marker that generally shows broad modifications ^2,4^,14, and we assessed this modification by NTU-CAT. A snapshot of the NTU-CAT peaks from six single blastocysts is shown in Fig. 6a, alongside the peaks from our previous ChIP-seq analysis of a cohort of blastocysts ^14^. The results showed that representative H3K27me3 modifications, such as those on *HOXA* and *PAX6* genes, could be detected using NTU-CAT. However, the H3K27me3 peaks obtained by NTU-CAT exhibited a lower resolution compared with those generated by ChIP-seq. In addition, inter-(ChIP-seq vs. NTU-CAT) and intra-(within the same method) assay correlations of the signals were lower compared with the case of H3K4me3 (Fig. 6b).

**Figure 6.**
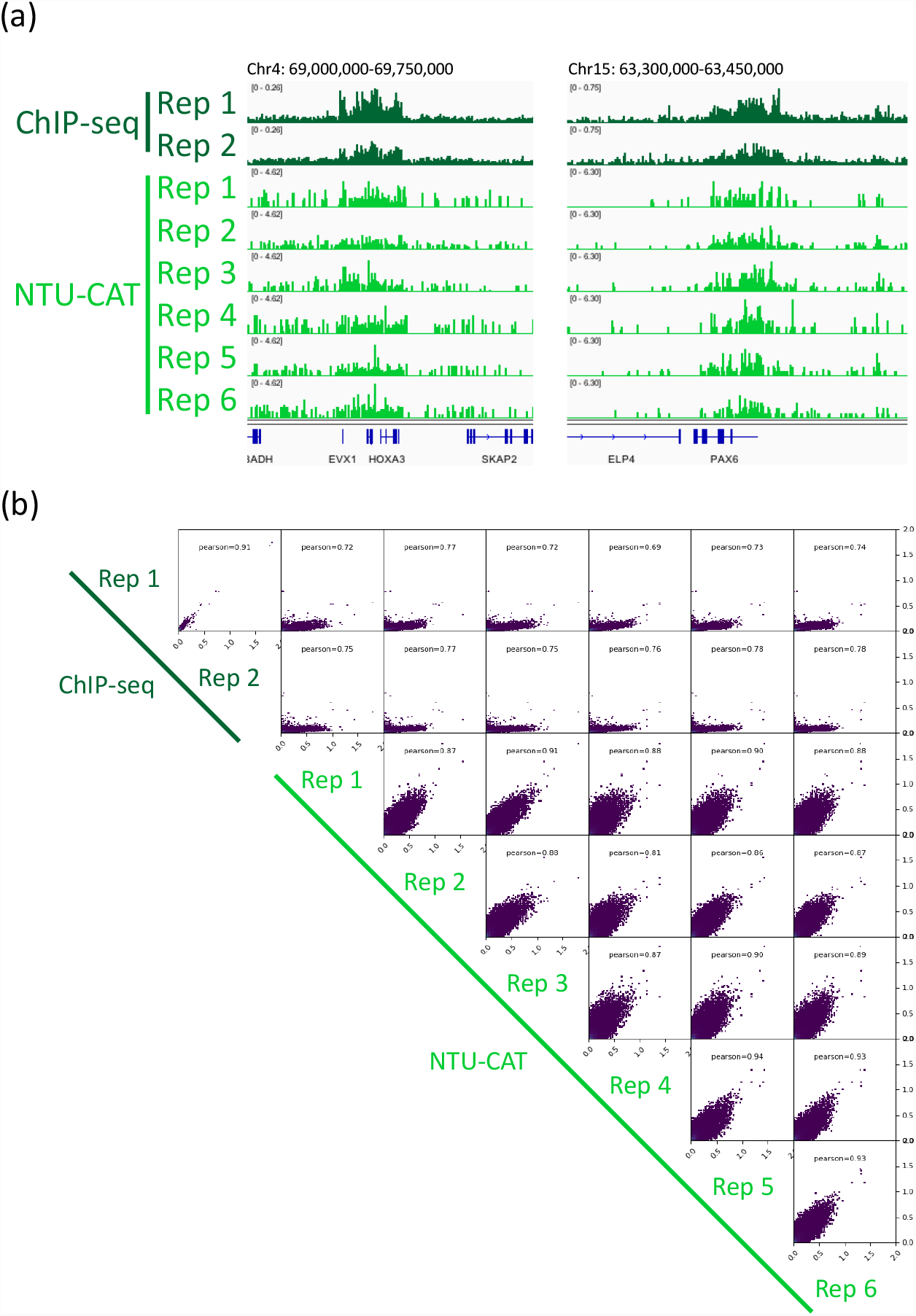
Comparison of the NTU-CAT and ChIP-seq results for H3K27me3 in bovine blastocysts. (a) Snapshot of the H3K27me3 landscape in representative positive regions, *HOXA* and *PAX6* genes, respectively. The NTU-CAT-derived peaks in six replicates were visualized using the Integrative Genomics Viewer ^19^ alongside the peaks from our previous ChIP-seq analysis of a cohort of blastocysts^14^. (b) Correlation analysis of duplicates of ChIP-seq and six replicates of NTU-CAT for H3K27me3 in bovine blastocysts. Scatterplots were generated using the same method as in Fig. 3.

### Typical characterization of single embryos by NTU-CAT-*XIST* activation

A promising advantage of single embryo analysis is the characterization of individual embryos. To explore the possibility of characterizing individual embryos based on a specific marker that can be detected by NTU-CAT, we focused on H3K4me3 modification of the *XIST* locus. The results showed a clear dichotomy between the presence or absence of H3K4me3 modification at this locus obtained by NTU-CAT (Fig. 7). This suggests that the NTU-CAT method can be applied to sexing of embryos using the presence or absence of *XIST* activation.

**Figure 7.**
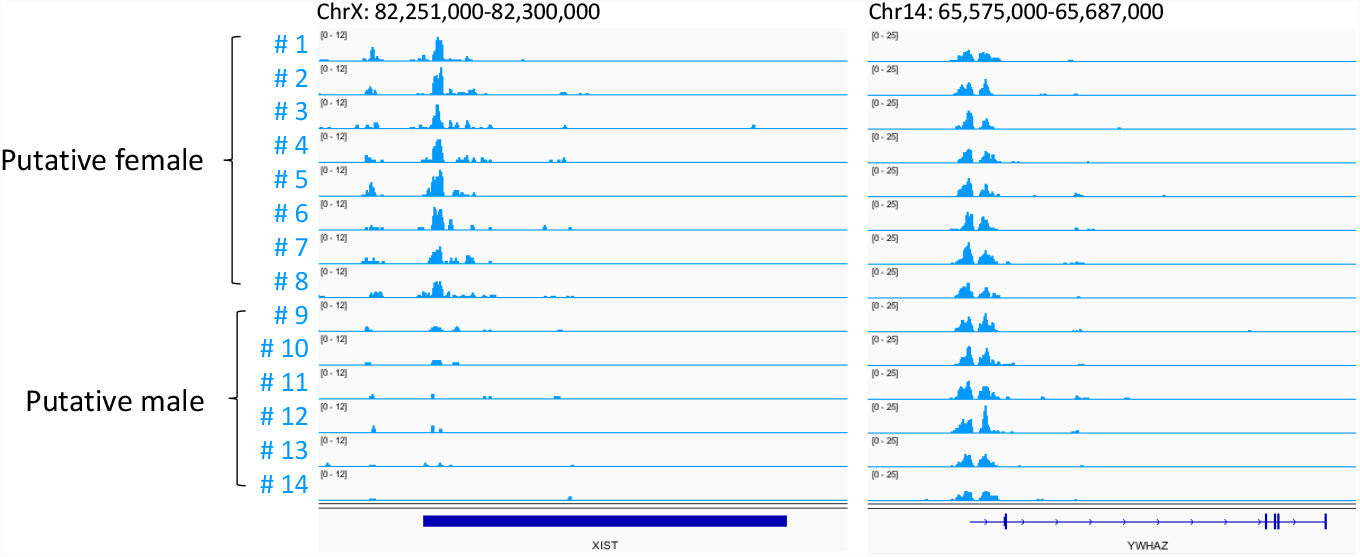
H3K4me3 modifications at the *XIST* locus in each embryo. The right panel shows an example of a gene with sex-independent H3K4me3 modifications, like many others, of *YWHAZ*.

## Discussion

Epigenomic analysis of single preimplantation embryos paves the way for the characterization of individual early embryos. In the present study, we further modified the CUT&Tag method ^9,10^, which can analyze histone modifications in a small number of cells, such that the embryo is handled in the absence of the solid-phase magnetic beads used for antibody and enzyme reactions in the original method. The original CUT&Tag approach utilizes concanavalin A-coated magnetic beads for fixing dispersed cells to the solid phase in order to facilitate handling and processing of the cells ^9,10^. However, preimplantation embryos are cell masses that can be transferred individually to any reaction solution in the experimental process using a fine pipette; we attempted to handle the embryos using this approach and named the method as NTU-CAT (Fig. 1).

As a result, despite some limitations, NTU-CAT generated genome-wide profiles of representative histone modifications, H3K4me3 and H3K27me3, even from single embryos, comparable to the results of the conventional ChIP-seq method using a cohort of embryos. For both modifications, we demonstrated the overall similarity of the signals detected between ChIP-seq and NTU-CAT (Figs. 2, 3, and 6).

For H3K4me3, some peaks were found to be uniquely more abundant in NTU-CAT (Fig. 2a), and the shape of the average profile of the peaks near the TSSs of genes differed between both methods such that the “valleys” detected by ChIP-seq were not observed with NTU-CAT (Fig. 2b). This may be due to the bias of Tn5 transposase to preferentially cut open chromatin regions ^12^. To estimate the magnitude of this bias, we evaluated the generation of false positive peaks by the comparison of NTU-CAT, ChIP-seq ^11^, and ATAC-seq peaks ^13^. As a result, it was found that there were 10%-15% false positive peaks per sample that were detected possibly due to the open chromatin bias (Fig. 5a). However, the remaining peaks were consistent between ChIP-seq and NTU-CAT, and the average area of false positive peaks were relatively smaller than those of total NTU-CAT peaks (Fig. 5a and b). Furthermore, high correlations of the signals per genomic bins were obtained between ChIP-seq and NTU-CAT (Pearson correlation = 0.87-0.91) and within the NTU-CAT replicates (0.95-0.98, Fig. 3).

For H3K27me3, the reproducibility between both methods (Pearson correlation = 0.69-0.78) and within the NTU-CAT replicates (0.81-0.94) was lower compared with the case of H3K4me3. However, NTU-CAT was able to detect the typical H3K27me3 peaks that can be detected by ChIP-seq, albeit at a lower resolution (Fig. 6a).

The single embryo-based analysis of histone modifications provides specific details about each embryo. For example, we could detect H3K4me3 modification at the *XIST* locus in about half of the examined embryos, suggesting that these embryos were female and exhibited *XIST* activation leading to X chromosome inactivation (Fig. 7).

In summary, despite NTU-CAT having some limitations that should be noted, such as false positive peaks and lower resolution for broad modifications, we concluded that it is a promising replacement for ChIP-seq with the great advantage of being able to analyze individual embryos.

## Methods

### *In vitro* production of bovine embryos

This study was approved by the Animal Research Committee of Kyoto University (permit number R3-10) and was carried out in accordance with the Regulation on Animal Experimentation at Kyoto University. The bovine ovaries used in this study were purchased from a commercial abattoir as by- products of meat processing, and the frozen bull semen used for *in vitro* fertilization (IVF) was also commercially available. *In vitro* production of bovine embryos by IVF was performed as previously described ^11,14^. Blastocyst-stage embryos at 170 h post IVF (day 7) were collected individually.

### NTU-CAT

The blastocysts were freed from the zona pellucida by using pronase and, from this point on, individual embryos were handled with separate solutions and pipettes to avoid mixing each other. The basal kit for CUT&Tag used was a CUT&Tag-IT Assay Kit (Active Motif). After washing the zona-free blastocysts with phosphate-buffered saline containing 0.01% (w/v) polyvinyl alcohol and 1% (v/v) Protease Inhibitor Cocktail (PIC), they were individually allocated to 50 μL Antibody Buffer with 1 µL (1.4 – 1.6 µg) of the target primary antibody, 0.05% (w/v) digitonin, and 1% (v/v) PIC in the wells of a round bottom-shaped 96-well plate. The blastocysts were incubated overnight at 4°C with gentle shaking (400 rpm). After the primary antibody reaction, the blastocysts were transferred to wells containing 100 µL Dig-Wash buffer with a secondary antibody (guinea pig anti-rabbit IgG antibody), 0.05% (w/v) digitonin, and 1% (v/v) PIC and incubated for 1 h at room temperature (400 rpm). After three washes with 180 µL Dig-Wash Buffer supplemented with digitonin and PIC in the 96-well plate, the blastocysts were transferred to wells containing 100 µL Dig-300 Buffer with pA-Tn5 Transposomes, 0.01% (w/v) digitonin, and 1% (v/v) PIC and incubated for 1 h at room temperature (400 rpm). After three washes with 180 µL Dig-300 Buffer supplemented with digitonin and PIC on the 96-well plate, the blastocysts were transferred to individual microcentrifuge tubes (Eppendorf 0030 108.051) containing 125 µL Tagmentation Buffer with 0.01% (v/v) digitonin and 1% (v/v) PIC and incubated for 1 h at 37°C without shaking.

After tagmentation, 4.2 μL of 0.5 M EDTA, 1.25 μL of 10% SDS and 1.1 μL of 10 mg/mL proteinase K were added to the tubes and incubated for 1 h at 55°C or overnight at 37°C with vigorous shaking (1,300 rpm). The tubes were heated at 70°C for 20 min to inactivate proteinase K, and then cooled to room temperature. SPRISelect beads (Beckman Coulter, 145 µL) were added to the tubes, vortexed for 1 min, and incubated for 10 min. The tubes were placed on a magnet stand for 4 min to collect the magnetic beads and the liquid was removed. The beads were washed twice with 1 mL of 80% ethanol. After drying the bead pellets for 2-5 minutes, 35 µL DNA Purification Elution Buffer was added and the tubes were vortexed and left to stand for 5 min. The tubes were placed on a magnet stand for 4 min to collect the magnetic beads and the liquid containing tagmented DNA was transferred to PCR tubes. PCR amplification of sequencing libraries was performed in a volume of 50 µL using 30 µL tagmented DNA and i7 and i5 indexing primers according to the manufacturer’s protocol. The PCR condition was as follows: 72°C for 5 min; 98°C for 30 s; 20 cycles of 98°C for 10 s and 63°C for 10 s; final extension at 72°C for 1 min; and hold at 10°C. Post-PCR library purification was performed with 55 µL SPRISelect beads (vortex 1 min, stand for 5 min, and bead collection 4 min) and 180 µL of 80% ethanol as described above. Finally, the sequencing libraries were eluted in 25 µL DNA Purification Elution Buffer.

### DNA sequencing and data processing

Sequencing was performed on a HiSeqX (Illumina) as paired-end 150-base reads. The sequencing reads were quality checked, merged, and aligned to the bovine genome (Bos_taurus_UMD_3.1.1/bosTau8, June 2014) using Bowtie 2 ^15^. Handling of sam and bam files were performed by using Samtools (http://www.htslib.org/). Mapping duplicates were removed by Picard (http://broadinstitute.github.io/picard/). The generated bam files were converted to bigWig (bw) files by using the bamCoverage tool of deepTools (https://deeptools.readthedocs.io/en/develop/) with counts per million normalization. The correlation plots between the experiments were made from the bw files fed to deepTools. The peaks were called using MACS ^16^. The annotation of called peaks to genomic regions was generated using CEAS ^17^. Average H3K4me3 signal profiles were generated by ngs.plot ^18^. Peaks were visualized using the Integrative Genomics Viewer (IGV) ^19^. The common and specific peaks between groups were identified using bedtools (https://bedtools.readthedocs.io/en/latest/) with the -wa and -v option, respectively to calculate the false positive rate (FPR).

### Publicly available data

The following raw data from publicly available databases were used: ChIP-seq of bovine blastocysts from our previous studies ^11,14^, i.e., rep1 and rep3 of GSE1612221 (H3K4me3) and rep1 and rep2 of GSE171701 (H3K27me3); RNA-seq of bovine blastocysts, Blastocyst_replicate1, 2, and 3 of GSE52415 ^20^; and ATAC-seq of bovine inner cell mass, ICM_rep1 of GSE143658 ^13^. For RNA-seq data, the three (blastocyst) datasets were merged and expression levels in RPKM values were calculated as previously described ^21^. The genes were evenly divided into three categories as high, medium, and low expression levels according to the calculated RPKM values.

## Acknowledgments

The authors deeply thank the staff at the Kyoto-Meat-Market for allowing us access to bovine ovaries. This work was supported in part by Livestock Promotional Subsidy from the Japan Racing Association and the Japan Society for the Promotion of Science (19H03104 to S.I. and 19H03136 to N.M.).

## Author Contributions

S.I., Y.H., and N.M. conceived the experiments. K.S. and S.I. performed bovine IVF and NTU-CAT library preparation and sequencing. K.S., S.I., H.Y., S.H., and N.M. analyzed the results. K.S. and S.I. drafted the manuscript. All authors discussed the results and approved the manuscript.

## Additional Information

### Supplementary information

Supplementary data are submitted along with the main manuscript.

**Supplementary Table S1**.

**Table S1.** Data generated by NTU-CAT of bovine blastocysts in this study

| Table S1. Data generated by NTU-CAT of bovine blastocysts in this study |  |  |  |  |
| --- | --- | --- | --- | --- |
| Replicate | Total reads | Total mapped reads (%) <sup>1</sup> | Uniquely mapped reads (%) | Reads after de-duplication <sup>2</sup> |
| H3K4me3_Rep1 | 309,069,780 | 290,859,627 (94.11) | 253,440,564 (82.00) | 1,517,565 |
| H3K4me3_Rep2 | 271,397,942 | 254,467,978 (93.76) | 226,414,835 (83.43) | 1,366,333 |
| H3K4me3_Rep3 | 256,869,432 | 240,626,890 (93.68) | 210,370,289 (81.90) | 1,028,471 |
| H3K4me3_Rep4 | 183,987,624 | 172,417,072 (93.71) | 152,025,702 (82.63) | 936,256 |
| H3K4me3_Rep5 | 188,117,734 | 175,622,708 (93.36) | 155,765,943 (82.80) | 978,502 |
| H3K4me3_Rep6 | 191,796,858 | 180,940,019 (94.34) | 159,421,116 (83.12) | 1,153,830 |
| H3K4me3_Rep7 | 182,234,060 | 172,129,980 (94.46) | 150,799,519 (82.75) | 886,026 |
| H3K4me3_Rep8 | 137,009,656 | 128,671,827 (93.91) | 109,848,249 (80.18) | 798,472 |
| H3K4me3_Rep9 | 172,778,388 | 161,698,547 (93.59) | 143,109,357 (82.83) | 1,258,822 |
| H3K27me3_Rep1 | 194,624,022 | 183,087,772 (94.07) | 126,286,155 (64.89) | 845,622 |
| H3K27me3_Rep2 | 196,073,114 | 184,942,697 (94.32) | 131,068,624 (66.85) | 1,555,708 |
| H3K27me3_Rep3 | 177,644,032 | 167,949,176 (94.54) | 123,051,701 (69.27) | 1,173,395 |
| H3K27me3_Rep4 | 294,872,164 | 281,759,640 (95.55) | 194,676,393 (66.02) | 793,021 |
| H3K27me3_Rep5 | 187,539,216 | 178,467,948 (95.16) | 128,827,707 (68.69) | 1,176,819 |
| H3K27me3_Rep6 | 371,009,066 | 353,574,975 (95.3) | 260,924,371 (70.33) | 1,081,875 |
| <sup>1</sup> Mapped using Bowtie 2. |  |  |  |  |
| <sup>2</sup> Processed by Samtools and Picard. |  |  |  |  |

### Data Availability

The NTU-CAT datasets have been deposited at Zenodo (https://zenodo.org/record/6002122).

